# A genome-wide assessment of the ancestral neural crest gene regulatory network

**DOI:** 10.1101/275875

**Authors:** Dorit Hockman, Vanessa Chong-Morrison, Daria Gavriouchkina, Stephen Green, Chris T. Amemiya, Jeramiah J. Smith, Marianne E. Bronner, Tatjana Sauka-Spengler

**Author notes:** Lead and corresponding author: Tatjana Sauka-Spengler.

## Abstract

The neural crest is an embryonic cell population that contributes to key vertebrate-specific features including the craniofacial skeleton and peripheral nervous system. Here we examine the transcriptional profiles and chromatin accessibility of neural crest cells in the basal sea lamprey, in order to gain insight into the ancestral state of the neural crest gene regulatory network (GRN) at the dawn of vertebrates. Transcriptome analyses reveal clusters of co-regulated genes during neural crest specification and migration that show high conservation across vertebrates for dynamic programmes like Wnt modulation during the epithelial to mesenchymal transition, but also reveal novel transcription factors and cell-adhesion molecules not previously implicated in neural crest migration. ATAC-seq analysis refines the location of known *cis*-regulatory elements at the *Hox-*α*2* locus and uncovers novel *cis*-regulatory elements for *Tfap2B* and *SoxE1*. Moreover, cross-species deployment of lamprey elements in zebrafish reveals that the lamprey *SoxE1* enhancer activity is deeply conserved, mediating homologous expression in jawed vertebrates. Together, our data provide new insight into the core elements of the GRN that are conserved to the base of the vertebrates, as well as expose elements that are unique to lampreys.

## Introduction

The neural crest (NC) is a migratory embryonic cell population that is unique to vertebrates and one of its defining features. Neural crest cells form in association with the developing central nervous system, which they delaminate from after undergoing an epithelial-to-mesenchymal transition (EMT). They subsequently migrate throughout the body to give rise to a plethora of tissue types including the neurons and glia of the peripheral nervous system, parts of the craniofacial skeleton and pigment cells of the skin^1^. The advent of the neural crest with its many tissue derivatives is thought to have played an essential role in the diversification of vertebrates^2,3^. Elucidating how the genetic signals involved in neural crest specification were modified over the course of vertebrate evolution is key to understanding how this diverse assemblage evolved and expanded^4^. This requires painting a detailed picture of how the neural crest gene regulatory network (NC GRN) functioned in the vertebrate ancestor. To this end, the sea lamprey, a basal vertebrate, serves as a good model based on its critical phylogenetic position. Morphologically these animals are considered “living fossils” with a body-plan that has remained consistent over at least the last 400 million years^5^.

The current view of the neural crest GRN has been compiled and refined from data generated in many jawed vertebrates^6^. By taking a candidate gene approach to compare lamprey and gnathostome transcription factors and signalling molecules, we previously showed that many key neural crest genes were conserved in expression and function between lamprey and jawed vertebrates^7^. These results suggested that the basic neural crest GRN was already present at the base of vertebrates, although some key regulators were missing from the lamprey neural crest specifier module^7^.

Recently, our understanding of the composition and function of the neural crest GRN in gnathostomes has been greatly increased with the advent of next generation sequencing techniques including RNA-seq, ChIP-seq and ATAC-seq^8–15^. Whereas the GRN of gnathostome neural crest has been greatly expanded, progress in reconstructing the neural crest GRN of jawless vertebrates has been limited due to incomplete genomic information. Recently, a germline genome assembly for the sea lamprey, that unlike previous assemblies^16^ is not affected by DNA-elimination^17,18^ and uses an integrated scaffolding approach to increase contiguity to near chromosome-scale resolution^19^, has made it possible to interrogate the regulatory genome of this basal vertebrate. Using this assembly, it is now possible to integrate gene expression data and *cis*-regulatory analyses on a genome-wide scale with increased confidence.

Here, we explore the dynamics of gene expression and chromatin accessibility during cranial neural crest specification and migration in the sea lamprey. By comparing our genome-wide representation of the active lamprey neural crest transcriptome to that of jawed vertebrates, our analyses highlight the components of the neural crest GRN that are conserved and therefore highly likely to be essential for neural crest specification. We analyse the chromatin accessibility in the neural crest cells of two different lamprey species, and find that cross-species mapping highights putative cis-regulatory elements. Importantly, we identify enhancer elements that drive expression in the premigratory and migratory neural crest of the lamprey, and provide evidence that regulation of a *SoxE* family gene is conserved between jawless and jawed vertebrates. By adapting high throughput tools to the lamprey, our data provide unique insight into the ancestral state of the neural crest GRN^20^.

## Results

### Dynamics of the developing neural crest transcriptome

As a first step, we obtained high quality cranial neural crest RNA-seq data at successive stages of development by dissecting the dorsal neural tube including premigratory, migrating and/ or post-migratory neural crest cells of the head at Tahara stage18 (T)18, T20 and T21 embryos (Fig.1a), respectively. In sea lamprey (*Petromyzon marinus*) embryos, neural crest cells reside within the neural folds which start to converge at T18 to form a neural rod and fuse at T20, when the first signs of neural crest migration have been reported^21,22^.

Reads were mapped to the sea lamprey germline genome assembly. A consensus transcriptome consisting of 120,207 transcripts at 72,171 genetic loci was assembled *de novo* (i.e. independent of current gene annotation) from the mapped dorsal neural tube datasets, combined with mapped RNA-seq datasets from whole heads and whole embryos at T20. 67,736 of the transcripts did not overlap with any annotated genes and thus represent candidate novel transcripts or transcribed transposable elements. The latter are present in large numbers in the lamprey genome but were not integrated in the current conservative gene model annotation that excluded repetitive elements^19^. Principal component analysis (PCA) of dorsal neural tube count data showed clear separation along principal component 1 (PC1), which accounted for 90% of the variance, reflecting the developmental stage at which the tissue was sampled (Fig. 1b). Both PCA and regression analysis confirmed that the biological replicate RNA-seq datasets at each stage were highly correlated, demonstrating high reproducibility (Supplementary Fig. 1). Differential expression analysis between the T18 and T21 samples, which represent the premigratory and migratory cranial neural crest respectively, revealed 9,106 differentially expressed genes (adjusted *p-value <*0.05). Of these, 5,400 were enriched at T21, while 3,706 were depleted (Fig. 1c).

**Figure 1:**
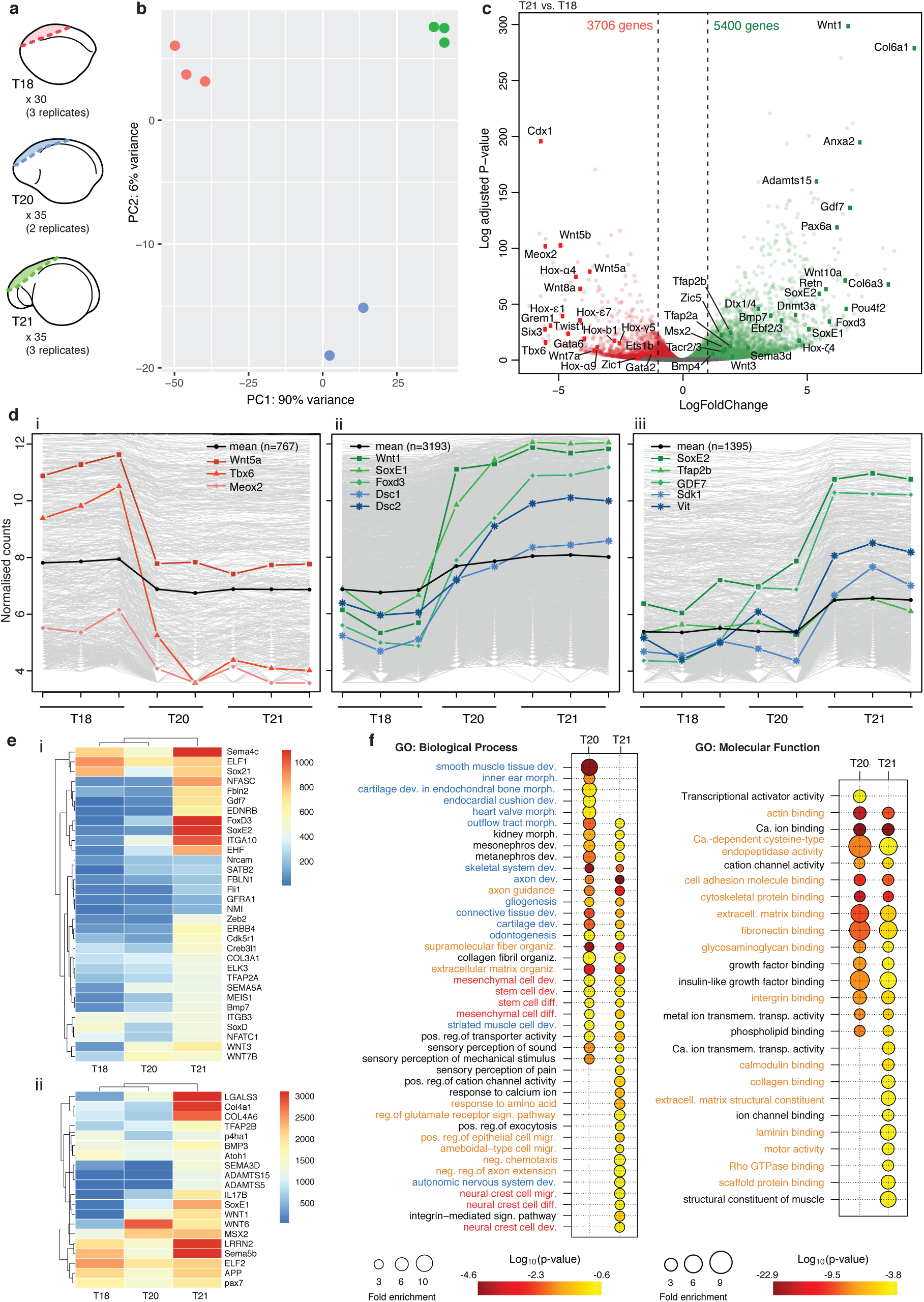
Dynamics of the d-eveloping neural crest gene expression profile. **a**, Schematic depicting the dorsal neural tube region dissected from T18, T20 and T21 lamprey embryos for RNA-seq. **b**, PCA of rlog-transformed gene expression count tables for 56,319 genes with non-zero read counts. PC1, which accounts for 90% of the variance is stage dependent (colours indicate stage as in a). **c**, Volcano plot of differential expression analysis between T21 and T18 (*p-value <*0.05; green, enriched; red depleted at T21). Coloured dots and labels indicate genes previously known to be enriched or depleted in the developing neural crest. *d*, Clusters of highly correlated genes identified by WGCNA (i, down-regulated after T18; ii, up-regulated at T20; iii, up-regulated at T21), showing genes that are known to be down-regulated (red) or up-regulated (green) in the neural crest, as well as up-regulated genes that have not been previously implicated in neural crest development (blue). **e**, Heatmaps of the average variance stabilised normalised gene counts for selected genes from WGCNA clusters 2 and 3, showing increased expression at T21. Low-level (i) and high-level (ii) expressing genes are shown. **f**, Bubble plots summarising enrichment and *p-values* for the most significant GO biological process and molecular function terms associated with enriched genes at T20 and T21 (only terms enriched more than three-fold are shown). The neural crest programme (red) is setup at T20, and continues at T21. Several GO terms associated with cell migration (orange) and neural crest derivatives (blue) are also present.

First, we assessed the dynamics of signalling molecules and transcription factors known to be expressed during neural crest specification making use of the germline genome annotation that used multispecies BLAST alignments to assign lamprey gene models to likely vertebrate homologues^19^. As expected, several *bona fide* neural crest markers were enriched at T21 when compared to T18 (Fig. 1c, Supplementary File1). For example, *Wnt1*, which plays a role in establishing the neural plate border and is maintained in the dorsal neural tube^23^, was one of the most significantly enriched genes as were *Wnt3* and *Wnt10*. In contrast, several *Wnt* homologues (*Wnt5a*/*b, Wnt7a, Wnt8a*) were depleted at T21, consistent with studies showing that *Wnt* expression is modulated during neural crest delamination and migration^14,24^, with a switch from canonical Wnt signalling critical for specification to involvement of Wnt/PCP pathway during cell migration^15,25,26^. Neural crest specifier genes like *SoxE* genes (*SoxE1* and *SoxE2*), *FoxD3, Msx2, Tfap2A* and *Tfap2B* (Fig. 1c) were increased by at least two fold at T21 whereas other genes including several *Hox, Tbx* and *Gata* transcription factors were depleted (Fig. 1c), analogous to previous observation in gnathostomes^8^. In contrast to jawed vertebrates^27,28^, both *Ets1b* and *Twist1* were depleted at T21, confirming our previous findings regarding their absence from the migratory neural crest of lamprey^29^.

Weighted Gene Co-expression Network Analysis (WGCNA)^30^ revealed 12 gene clusters with significantly higher gene expression at T18, and 13 gene clusters with significantly higher gene expression at T21, mirroring the results from our differential expression analysis (Supplementary File2, Supplementary Fig. 2). This approach delineated patterns of all genes expressed in neural crest cells. For example, *Tbx6* and *Wnt5a* were placed in a cluster of 767 genes with an expression trend that showed a drop from T18 to T20, and remained low at T21 (Fig. 1di; Supplementary File2:cluster1). The largest cluster (3,193 genes) showed an increase in expression from T18 to T20, maintained at T21. This contained key ‘neural crest specification module’ genes such as *SoxE1, Foxd3, Wnt1, Pax7, Msx2* and *Tfap2A* (Fig. 1dii, 1e; Supplementary File2:cluster2). Interestingly, these transcription factors were co-expressed with cell adhesion and cytoskeletal factors known to be involved in neural crest emigration (*Integrin*[*ITG*]*A2* /*A10* /*B2, Galectin-3* [*Lgals3*], *Interleukin*[*IL*]*17*, etc.). Several ‘neural crest migration module genes’, including *SoxE2, Tfap2B* and *Gdf7*, were placed in the next largest cluster (1,395 genes), which displayed low expression at both T18 and T20, increasing at T21 (Fig. 1diii, 1e; Supplementary File2:cluster3). Other co-regulated transcription factors involved in neural crest migration were also placed in this cluster (such as *Sox21* and *Zeb2*), as well as signalling receptors and ligands (*ERBB4, Ednrb, Sema3D* /*4C* /*5B*), secreted matrix remodelling enzymes (*MMP13, ADAM10, ADAMTS15*), collagens (*Col3a1* /*4a1* /*4a6*), and other lamprey orthologues involved in organisation of the extracellular matrix (*Prolyl 4-hydroxylase subunit alpha-1* [*P4ha1*], *Fibulin* [*Fbln2*] and *Creb3l1*) (Fig. 1e). This cluster also featured downstream effectors ensuring proper differentiation into neural crest derivatives, such as melanocytes (*RAB32, Sox21*), neurons (*Nrcam, Atoh1, Netrin, Neurofascin* orthologues, *LRRN2*) and glia (*SoxD, GFRA1, Cdk5r1, APP* orthologues)(Fig. 1e).

**Figure 2:**
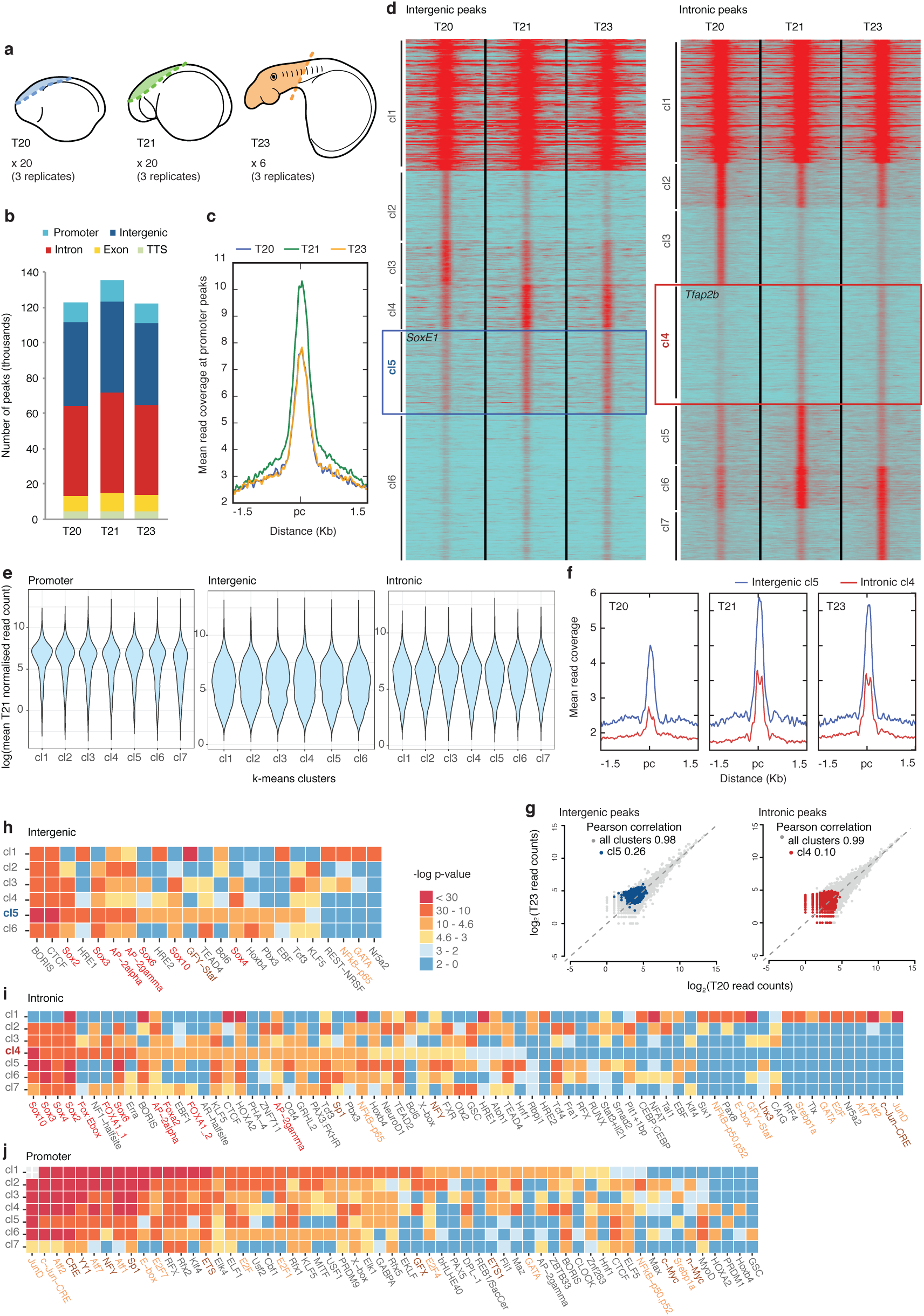
Profiling of chromatin dynamics in the developing neural crest reveals putative *cis*-regulatory elements involved in EMT. **a**, Schematics indicating the region of dorsal neural tube or head dissected from T20, T21 and T23 lamprey embryos for ATAC-seq. **b**, Genomic functional annotation of our ATAC-seq peaksets for all stages. **c**, Mean ATAC-seq read coverage map at each stage over our consensus promoter peakset (i.e. peaks associated with T21 enriched genes), showing higher read coverage at T21. **d**, Heatmaps depicting k-means linear enrichment clustering of ATAC-seq reads at all stages across consensus intergenic and intronic peaksets. Boxes indicated the large “EMT” clusters that show enriched signal at T21 and T23. **e**, Violin plots visualising the distribution of mean normalised T21 read counts for genes *k*-means clusters. Gene expression associated with promoter peak clusters (annotated and novel promoters) is higher and less variable than that for genes associated with intergenic and intronic clusters. **f**, Mean ATAC-seq read coverage maps at each stage for “EMT” clusters (intergenic cluster 5 in blue; intronic cluster 4 in red), showing higher coverage at T21 and T23. **g**, Scatterplot between T20 and T23 ATAC-seq read counts over consensus intergenic and intronic peaksets. Pearson correlation coefficients (r) for all clusters (grey) and for “EMT” clusters (intergenic cluster 5 in blue; intronic cluster 4 in red) are given. **h-j**, Transcription factor binding motif enrichment analysis for intergenic (h), intronic (i) and promoter (j, annotated and novel promoters) *k*-means clusters. Neural crest master regulator motifs are highlighted in red. Motifs shared between intergenic and intronic cluster 1 and promoter clusters are highlighted in orange. Canonical promoter motifs are highlighted in brown. Pc; peak centre.

Importantly, the two largest WGCNA clusters contained genes that have not previously been implicated in neural crest development, including genes coding for several cell-adhesion molecules, such as desmocolins (*Dsc1* /*2*) and *Sdk1*, and known extracellular matrix proteins, such as *vitrin* (Fig. 1dii,diii). Many novel transcription factors, as well as those that have been shown to play a role much later in neural crest development, were also placed in these clusters (Fig. 1e; Supplementary File2). For example, Nmi (N-myc interactor), known to interact directly with Sox10^31^ and inhibit canonical Wnt signalling in cancer cell lines^32^, showed increased expression at T20, while EHF (Ets homologous factor, also known as Epithelial Specific Ets-3), proposed to play a role as a tumour-suppressor in prostate cancer^33^ and oncogene in ovarian cancer^34^, showed greatly elevated expression at T21. *Fli1* and *Satb2*, which are both known to be expressed in the developing branchial arch cartilage and mesenchyme^35–37^, and Nfatc, which has been shown to form a complex with Sox10 during Schwann cell differentiation^38^, were also elevated at T21.

Gene Ontology (GO) analysis of genes enriched at T20 and T21 revealed overrepresentation for terms associated with early neural crest specification, cell migration and later neural crest derivatives (Fig. 1f). GO Terms associated with processes that have not been previously implicated in neural crest development were also present. These include glutamate signalling, organ (heart and kidney) morphogenesis and sensory perception of pain, sound and mechanical stimuli. Thus, our dataset captured the gene expression dynamics involved in cranial neural crest cell migration and delamination, while also providing insight into novel pathways that may be specific to lamprey neural crest development.

Taken together, our RNA-seq data confirms, with a higher level of detail, previous findings that a large proportion of the neural crest GRN is conserved to the base of the vertebrate family tree^2,29^. Importantly, our analyses also reveal many novel factors, whose role in neural crest development and diversification warrants further investigation.

### Genome-wide assessment of chromatin accessibility by ATAC-seq

With full transcriptome data in hand, we next sought to explore the regulatory connections between players in the neural crest GRN. To this end, it is essential to identify *cis*-regulatory elements that control gene expression specifically in the neural crest. ATAC-seq reveals regions of accessible chromatin, and thus enables a genome-wide assessment of the locations of putative *cis*-regulatory elements^39^. We used ATAC-seq to analyse chromatin accessibility in dissociated cells isolated from sea lamprey dorsal cranial neural tubes or whole heads at T20, T21 and T23 (Fig. 2a), representing the migratory and post-migratory neural crest. ATAC-seq datasets were mapped to and analysed within the context of the sea lamprey germline genome assembly^19^. Mapped ATAC-seq biological replicate datasets were highly correlated (Supplementary Fig. 3) and the insert size distribution of the mapped ATAC-seq libraries showed a stereotypical 150 bp periodicity (Supplementary Fig. 4a) which is consistent with the nucleosome occupancy of chromatin^39^ and demonstrates the quality of the ATAC-seq experiments.

**Figure 4:**
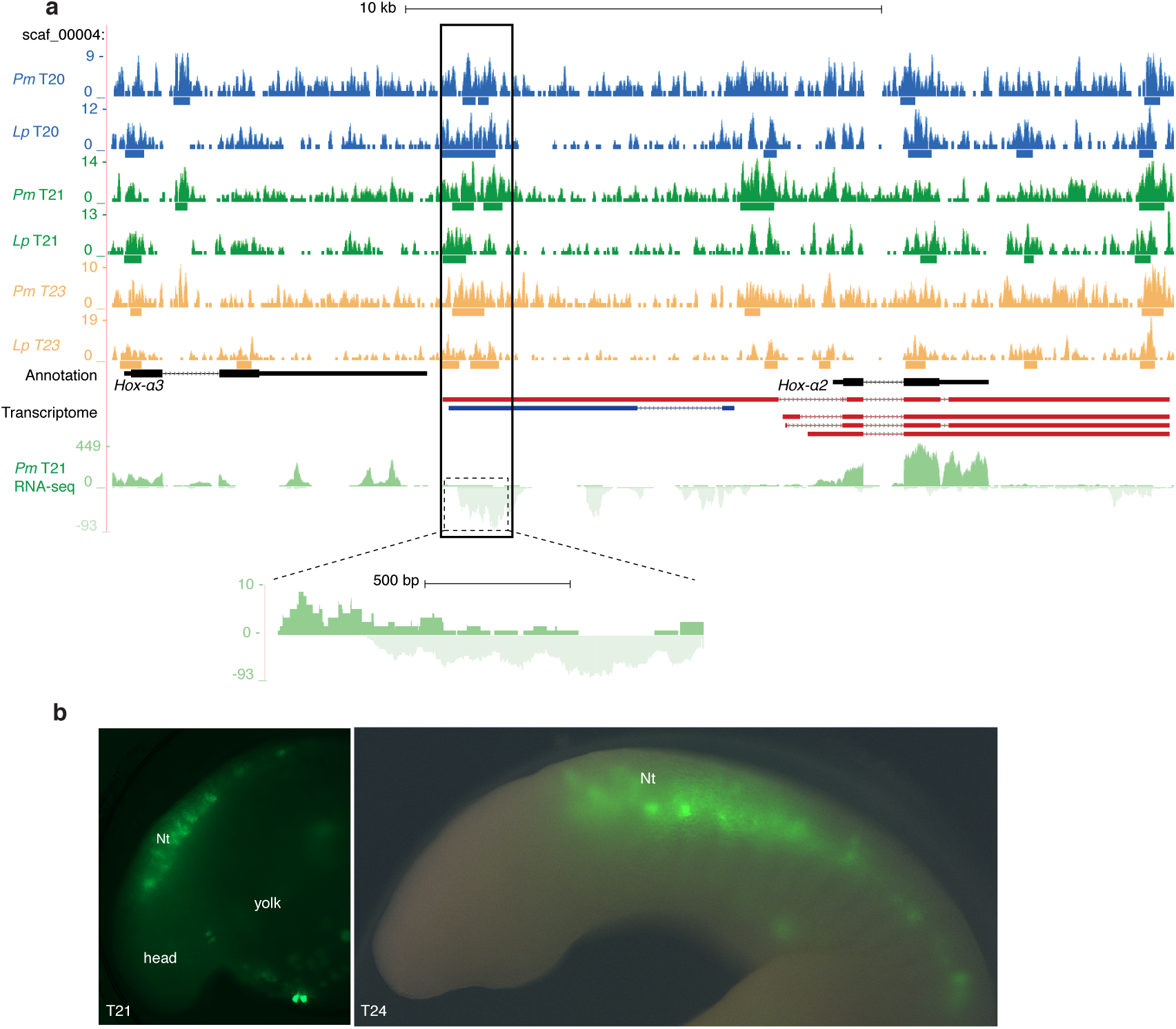
Characterisation of a *Hox-* α*2* enhancer and associated transcription. **a**, The *Hox-*α*3* /*Hox-*α***2*** locus in the sea lamprey germline genome, with representative ATAC-seq coverage tracks from the sea lamprey (Pm) and brook lamprey (Lp) for each developmental stage, as well as RNA-seq coverage tracks from a representative T21 sample indicating directional transcription. Bars below ATAC-seq coverage plots indicate peak regions. The black box indicates the region tested in enhancer-reporter assays, and the dashed box highlights bidirectional transcription over this region. *De novo* assembled transcripts for the *Hox-*α***2*** locus are shown maroon (sense) and dark blue (anti-sense). **b**, GFP reporter expression in lamprey embryos injected with the *Hox-*α***2*** enhancer-reporter construct at 1-cell stage and allowed to grow to indicated stages. GFP reporter expression is seen in the neural tube.

**Figure 3:**
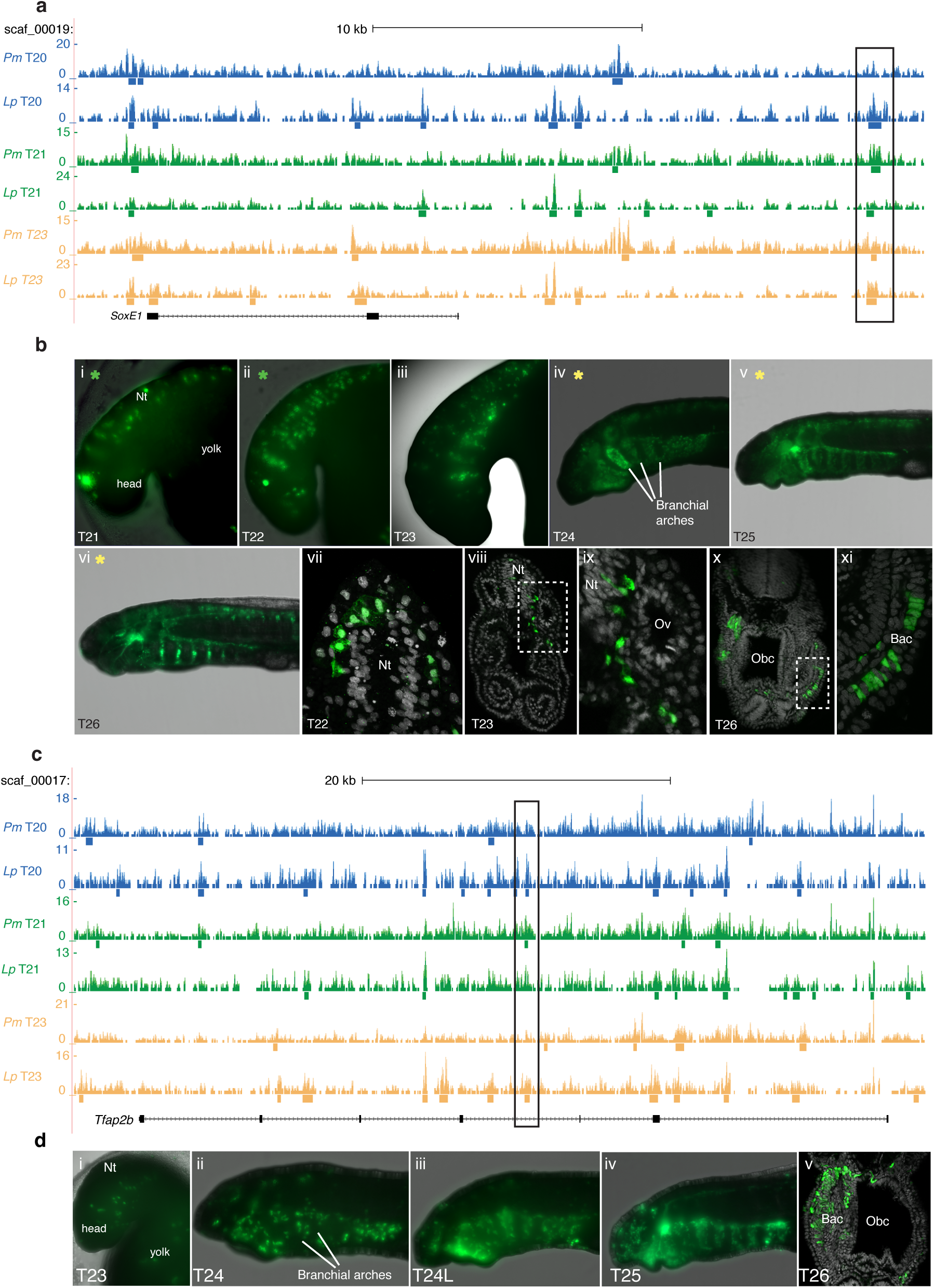
Tissue-specific enhancer activity in the lamprey neural crest. **a**, and **c**, The *SoxE1* (a) and *Tfap2B* (c) loci of the sea lamprey germline genome, with representative ATAC-seq coverage tracks from the sea lamprey (Pm) and brook lamprey (Lp) for each developmental stage. Bars below coverage plots indicate peak regions. The black box indicates the region tested in enhancer-reporter assays. **b**, GFP reporter expression in lamprey embryos injected with the *SoxE1* enhancer-reporter construct at the 1-cell stage and allowed to grow to indicated stages. **i-vi** are whole mount views, **vii-xi** are sections showing GFP^+^ cells delaminating from the neural tube (vii) migrating between the neural tube and otic vesicle (viii-ix) and contributing to the branchial arch cartilage (x-xi). **d**, GFP reporter expression in lamprey embryos injected with the *Tfap2B* enhancer-reporter construct at 1-cell stage and allowed to grow to indicated stages. i-iv are whole mount views, v is a transverse section showing GFP^+^ cells in the branchial arch cartilage. Coloured stars indicate panels showing the same embryo at successive developmental stages. Dashed boxed regions indicate regions magnified in adjacent panels. Bac, branchial arch cartilage; Obc, orobranchial cavity; L, late; Nt, neural tube; Obc, orobranchial cavity; Ov, otic vesicle.

Peak detection and annotation, using the *de novo* assembled consensus transcriptome generated in this study as a reference, was consistent across stages, with the majority of peaks found in intergenic and intronic regions where *cis*-regulatory elements are expected to be located (Fig. 2b). To focus our analyses on peaks associated with neural crest GRN genes, consensus peaksets for each annotation category (promoter, intergenic and intronic) were filtered to only contain peaks that were associated with genes enriched at T21. Promoter peak groups were further filtered to only contain elements associated with the promoters of genes annotated in the sea lamprey germline genome assembly (i.e. overlapping with a region up to 2 kb upstream from annotated gene models).

*K*-means clustering of the ATAC-seq signal over our consensus peaksets (8,998 intergenic peaks; 17,908 intronic peaks; 1,860 promoter peaks) revealed the dynamics of chromatin accessibility genome-wide at neural crest gene promoters and putative *cis*-regulatory elements over the course of development (Fig. 2c-g). The ATAC-seq signal associated with the promoter peaks was highest at T21 for all clusters showing, as expected, that enriched gene expression correlated with increased promoter accessibility (Fig. 2c). Additionally, when all promoter peaks were taken into consideration (i.e. 10,286 annotated and novel promoter peaks), the expression level of genes associated with the promoter peaks was higher and less variable than that associated with the intergenic and intronic peak clusters (Fig. 2e, Supplementary Fig. 4b).

We were particularly interested in clusters containing *cis*-regulatory elements that play a role in regulating gene expression during the epithelial-to-mesenchymal transition (EMT) that initiates cranial neural crest migration. *K*-means clustering of intergenic and intronic peaks revealed two large clusters (intergenic cluster 5; intronic cluster 4) that displayed increased accessibility at T21 and T23 compared to T20 (Fig. 2d,f). Gene ontology terms associated with intergenic cluster 5 included ‘regulation of localisation’, ‘positive regulation of cell-substrate adhesion’, and ‘regulation of cell motility’, while terms associated with intronic cluster 4 included ‘cell-cell junction organisation’, ‘cytoskeleton reorganization’ and ‘positive regulation of cell migration’ (Supplementary Fig. 4c). Additionally these clusters contained elements associated with known neural crest GRN transcription factors, *SoxE1* (intergenic cluster5) and *Tfap2B* (intronic cluster4).

To quantify the significance of these “EMT” clusters, we plotted the ATAC-seq signal levels of our peaksets at T20 against those at T23 and calculated the Pearson correlation coefficient for all intergenic and intronic clusters (Fig. 2g). This analysis of correlation coefficients revealed that both “EMT” clusters were significantly offset from all other identified groups of accessible elements, suggesting that the dynamics of opening of putative *cis*-regulatory elements may single them out as potentially functional enhancers during neural crest development.

We next used transcription factor binding site motif analysis to further interrogate the ATAC-seq *k*-means clusters. Intergenic cluster 5 was significantly enriched for Sox and Tfap2 binding sites, while intronic cluster 4 displayed a similar profile with the addition of Fox transcription factor binding sites (Fig. 2h-i). The presence of binding motifs for key neural crest transcription factors further suggested that these clusters very likely harbour *cis*-regulatory elements that provide connections between neural crest GRN players. In addition, enrichment of CTCF binding sites in all intergenic clusters further suggests these peaks may represent putative *cis*-regulatory elements^40^. Cluster 1 for both intergenic and intronic peaksets had a largely distinct TF binding site profile from the other clusters, which more closely resembled the binding profile of our promoter peakset (annotated and novel promoter peaks) (Fig. 2j), and was enriched for motifs found in the HOMER promoter motif library. These clusters also consisted of peaks that displayed a broad, open profile at all analysed stages (see Fig. 2d). Therefore, it is likely that cluster 1 for both intergenic and intronic peaksets represent peaks that show promoter-like activity.

### Identification of active neural crest-specific cis-regulatory elements

To test whether our ATAC-seq peaksets harboured active neural crest-specific *cis*-regulatory elements, we chose peaks from our intergenic and intronic“EMT” clusters that were associated with loci of known neural crest GRN genes to use in enhancer-reporter expression assays. A region of ~1.5 kb encompassing the ATAC-seq positive accessible chromatin region to be tested was cloned into the HLC reporter vector for transient transgenesis in 1-cell stage lamprey embryos^41^. An element, located 16.6 kb downstream of the *SoxE1* gene locus (Fig. 3a), drove highly specific reporter expression in the delaminating neural crest cells from T21 (observed in 195 out of 1,337 injected embryos) and labelled the cells as they migrated into the branchial arches and contributed to known neural crest-derived structures, such as the branchial arch cartilage (Fig. 3b). Similarly, an element located in the third intron of the *Tfap2B* gene drove reporter expression in the migrating neural crest from T23 (observed in 25 out of 340 injected embryos) and labelled neural crest derivatives at later stages (Fig. 3c-d). The location of both of these *cis*-regulatory elements overlapped with peaks in similar ATAC-seq datasets that were collected from the brook lamprey (*Lampetra planeri*) and mapped to the sea lamprey germline genome assembly (Fig. 3a,c).

This surprising finding suggests conservation of *cis*-regulatory elements across different lamprey species, which were separated at least 40 MYA^42^. Our analysis thus suggests a high degree of sequence conservation at the level of the functional non-coding regions of the genome of these two species, thus facilitating the identification of *cis*-regulatory elements using cross-species whole genome alignment of ATAC-seq data.

### Identification of a putative lncRNA associated with a lamprey Hox-α2 enhancer

Our ATAC-seq dataset can also be used to refine known *cis*-regulatory regions into modules that drive expression in specific tissues. Recently, sea lamprey *cis*-regulatory elements that drive gene expression in the developing neural tube, somites and neural crest were identified within a 9-kb region upstream of the *Hox-*α*2* locus, while elements that drive expression in the neural crest and somites alone were located within 4kb of the *Hox-*α*2* locus^43^. This was confirmed in our ATAC-seq dataset. We found that a *∼* 1.5kb region encompassing an ATAC-seq positive element at *∼* −8.5 kb drove reporter expression that was restricted to the neural tube (Fig. 4a-b).

Interestingly, within this locus, our RNA-seq data revealed bidirectional transcription, known to occur at active enhancers^44,45^ (Fig. 4a, bottom panel). Two novel transcripts from our transcriptome overlapped this region: a 12,770 bp sense transcript (Fig. 4a, maroon label) and a 4,206 bp, spliced antisense transcript (Fig. 4a, blue label). The longer sense transcript resembles the multiexonic enhancer (me)RNA transcript, similar to those reported in association with the ethryroid-specific intergenic enhancer, R4, located upstream of the *Nprl3* locus in mice^46^. As is the case with the R4 meRNA, the *Hox-*α*2* upstream enhancer appears to initiate the transcription of a unique alternative first exon, which is spliced onto an adjacent annotated exon and reads through the remaining exons of the *Hox-*α*2* gene. This results in a spliced transcript reminiscent of the annotated version, albeit with an extended 5^*t*^UTR or first exon. Therefore this enhancer, located ~8.5kb upstream of the *Hox-*α*2* locus, may be acting as alternative promoter (indeed, the peak was annotated as a promoter in our analyses). Alternatively, the production of this transcript might be a byproduct^46^, or perhaps a facilitator, of chromatin looping linking the upstream enhancer to the *Hox-*α*2* promoter.

The antisense transcript is one of 6,257 putative long non-coding (lnc)RNAs identified in our transcriptome (see Methods). 48% of these overlap with predicted lncRNA from adult sea lamprey brain, heart, kidney, and gonad RNA-seq datasets^19^, while 70% were associated with ATAC-seq positive regions. The gnathostome *HoxA* locus is known to harbour lncRNAs, including HOTAIRM1^47^ and HOTTIP^48^, both of which have been shown to modulate gene expression in *cis*. The putative lncRNA, identified between *Hox-*α*3* and *Hox-*α*2*, is significantly differentially expressed between T18 and T21 in the dorsal neural tube (4.7 fold change; *p.adj.*= 9.8E-21), suggesting it may play a role in regulating *Hoxa* gene expression during neural crest development.

### Lamprey SoxE1 enhancer activity is conserved in gnathostomes

One of the main aims of our study is to define the core components of the neural crest GRN that are conserved across vertebrates. This includes assessing whether the activity of neural crest enhancer elements present in a basal jawless vertebrate is conserved in jawed vertebrates, despite 500 million years of independent evolution^42^. To this end, we generated transgenic zebrafish carrying the lamprey SoxE1 enhancer upstream of a minimal promoter and GFP using the Activator (Ac)/Dissociation (Ds) (Ac/Ds) transposition system^49^. This system that facilitates highly efficient transgenesis in zebrafish resulted in a minimum of 7 independent integrations of the *SoxE1* enhancer:GFP cassette into the zebrafish genome, as determined by splinkerette PCR. While only weak reporter expression was visible in F_0_ embryos, the F_1_ generation displayed striking heterospecific reporter expression in the branchial arches by *∼* 30 hpf, closely mirroring the enhancer activity in the lamprey at T23 (Fig. 5ai-i’, Supplementary Fig. 5a). In older embryos (*∼* 60 hpf), GFP reporter expression was visible in several structures in the head that are known to receive neural crest contributions including the branchial arch cartilages and cranial ganglia, as well as developing and mature melanocytes, and putative Schwann cells in the trunk (Fig. 5aii-iii’).

**Figure 5:**
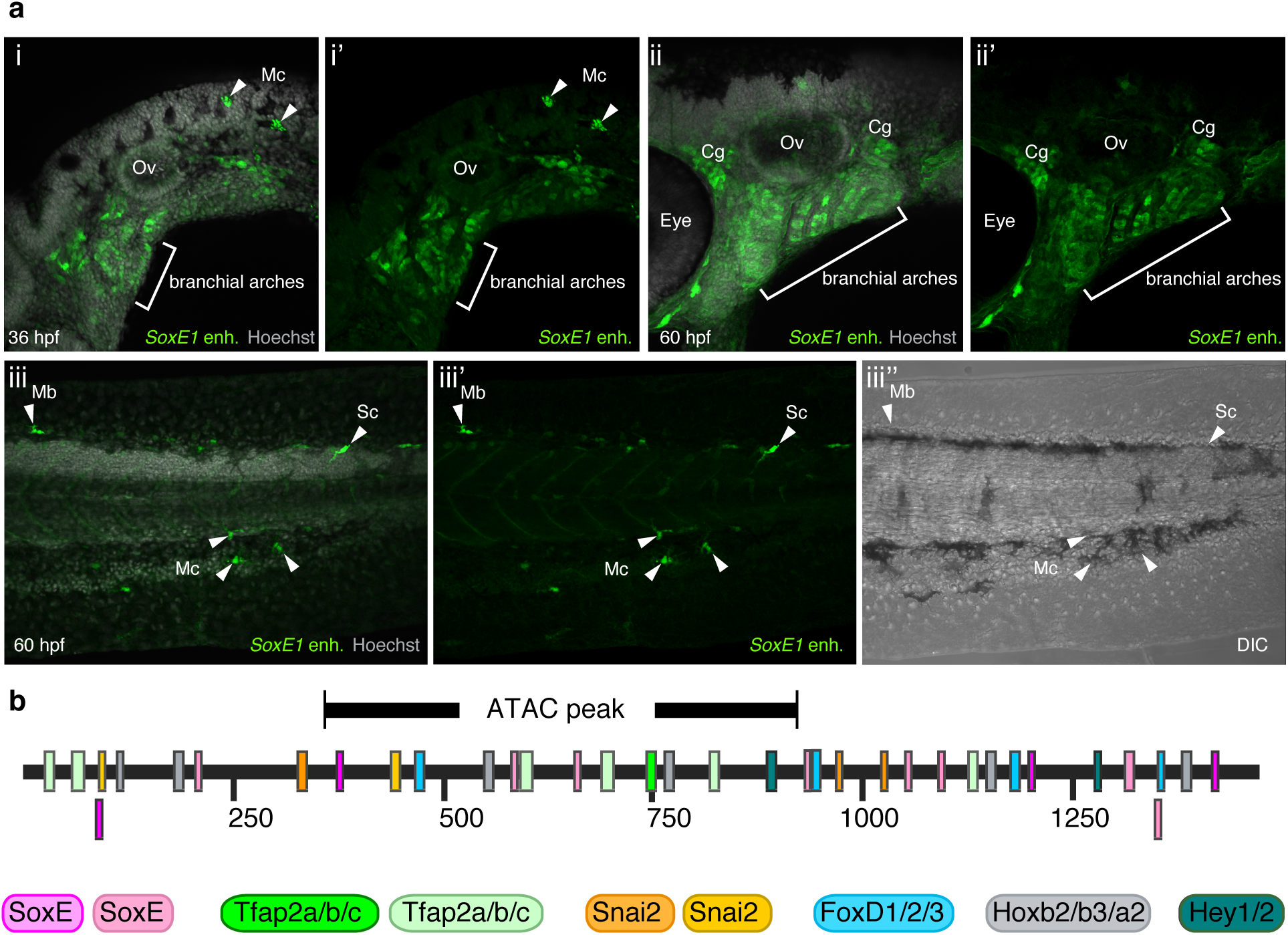
The activity of the lamprey *SoxE1* enhancer is conserved in gnathostomes. **a**, In a 36 hpf transgenic zebrafish GFP reporter expression is visible in the developing branchial arches and melanocytes (i-i’). At 60 hpf, GFP+ cells populate the branchial arch cartilage and cranial ganglia in the head (ii-ii’), while GFP+ melanocytes, melanoblasts and a putative Schwann cell are visible in the trunk (iii-iii”). **b**, Schematic of the lamprey *SoxE1* enhancer sequence with putative transcription factor binding motifs indicated by coloured boxes. Bright pink SoxE, bright green Tfap2A/B/C and bright orange Snai2 boxes indicate conserved canonical binding sites. The location of the ATAC-seq peak is indicated. Mb, melanoblasts; Mc, melanocytes; Ov, otic vesicle Sc, putative Schwann cell.

The fact that the lamprey *SoxE1* enhancer was active in the developing neural crest in zebrafish indicated that the transcription factor binding code characterised by our genome-wide motif analysis (Fig. 2h-i) and present in the lamprey *SoxE1* enhancer can be recognised by gnathostome transcription factors despite a lack of overt sequence conservation that would identify a homologous regulatory element. To test this hypothesis, we searched for the canonical binding sites of known neural crest transcription factors in the lamprey *SoxE1* enhancer. We found that the lamprey *SoxE1* enhancer harbours binding sites for several key neural crest transcription factors, including SoxE, Tfap2A/B/C and Snai2, but also for factors such as HoxA2 and HoxB3, which would be expected to restrict the enhancer activity in the cranial region to the hindbrain neural crest streams (Fig. 5b, Supplementary Fig. 5b). Rather than relying on extensive sequence conservation, our findings support the notion that the combination of short transcription factor binding site motifs are able to convey enhancer function^50^ and mediate the regulatory activity across evolutionary time.

Together, our cross-species enhancer-reporter analysis and transcription factor binding site search indicate that the regulation of *SoxE* gene expression in the migratory neural crest has been conserved from the base of the vertebrate tree and that enhancers with such conserved activity reflect the central lynchpin mediating the conservation of the neural crest GRN. Importantly, this suggests that regulation of one of the central players in the neural crest GRN has remained constant since the existence of the last common ancestor between jawed and jawless vertebrates.

## Discussion

Here, we present the most complete assembly to date of the lamprey neural crest gene regulatory network that can be directly compared with that of gnathostomes. Moreover, our combination of RNA-seq and ATAC-seq datasets in the developing lamprey neural crest provides a valuable platform for analysis of genome-wide gene expression and chromatin dynamics. The data not only enable identification of tissue-specific enhancers whose activity (but not overall sequence) was evolutionary conserved over several hundred million years, but also reveal putative non-coding RNA species whose functions remain to be explored in any vertebrate.

Analysis of the lamprey neural crest transcriptional network provides global insight into the evolution of neural crest transcriptional programmes. At premigratory neural crest stages (T20) in the lamprey, we observed similar gene enrichments in categories equivalent to those observed in zebrafish^15^. Our results reveal that the lamprey premigratory neural crest shows genetic signatures and significant functional enrichment in categories associated with a wide variety of both mesenchymal (smooth muscle, connective tissue, cartilage and tooth development) and neuronal (axonogenesis, gliogenesis) neural crest derivative fates. As development progresses and *bona fide* neural crest cells begin to delaminate (T21), enrichment terms changed to those characterising neural crest and stem cell programmes (neural crest and stem cell development, neural crest migration and differentiation) as well as autonomic nervous system formation. Interestingly, the neural crest GRN of lamprey differs from that of zebrafish in that it lacks the neural crest sub-programme involved in the specification of the enteric nervous system (ENS). This is consistent with studies showing that the lamprey may lack vagal neural crest and that the lamprey ENS may have much later onset and possibly different cell of origin^51^.

Analysis of co-expression clusters using WGCNA increases the resolution of the putative lamprey neural crest GRN^29^ and suggests links between early specification factors and their downstream effectors. Transcription factors and signalling molecules that are associated with the ‘neural crest specification module’ (e.g. *Wnt1, Msx2, Pax7, FoxD3, Tfap2A, SoxE1*) are upregulated at T20 and co-regulated with genes associated with the process of delamination, including cell adhesion and cytoskeletal factors. Later, at T21, transcription factors that are associated with neural crest migration (e.g *Sox21, Zeb2, Tfap2B*) are co-expressed with genes associated with active migration, including signalling receptors, their ligands and secreted matrix remodelling enzymes. Interestingly, these genes are also co-expressed with genes involved in the process of differentiation of neural crest derivatives, including neurons, melanocytes and glia. Future analysis using this type of classification as a resource to investigate direct links between co-regulated genes will allow expansion of the global structure of the lamprey neural crest GRN.

Our analysis of dynamic changes of chromatin accessibility in lamprey neural crest using ATAC-seq has enabled identification of developmentally regulated tissue-specific enhancers. ATAC-seq is a powerful tool for locating active *cis*-regulatory elements in pure cell populations^39^, different cell types^52^ and disease states^53^. Similar to studies from *C. elegans*, where ATAC-seq analysis of whole animals at early embryonic, larval and adult stages revealed dynamically regulated *cis*-regulatory regions with tissue specific activity^54^, we have performed this analysis using dissected cell populations predominantly, but not exclusively, comprised of neural crest cells at three successive stages of lamprey embryonic development. Together, these studies show that the dynamic signature associated with changes in chromatin accessibility over time can be used to pinpoint putative tissue-specific regulatory regions. Transcription factor binding site motif analysis provides further support for a neural crest-signature in our ATAC-seq peak clusters as the binding motifs for several key neural crest transcription factors were found to be enriched. This analysis also suggests that our dataset likely contains diverse *cis*-regulatory elements, including insulators (CTCF binding sites), promoters and enhancers that are being dynamically regulated during development.

Interestingly, we show that heterospecific analysis of ATAC-seq data from different lamprey species can provide clues for the location of conserved *cis*-regulatory regions. Although the evolutionary distance between the sea lamprey and gnathostomes precludes identification of *cis*-regulatory elements based on sequence conservation, the ATAC-seq reads from brook lamprey, *L.planeri*, were successfully mapped cross-species to the genome of the sea lamprey, *P. marinus*. This suggests that the putative functional non-coding elements have been conserved between the two lamprey species over the last 40 million years^42^. Thus, mapping the brook lamprey ATAC-seq data to the sea lamprey genome has enabled identification of conserved genomic regions (i.e. putative *cis*-regulatory elements).

Despite a lack of overt sequence conservation between the non-coding regions of the lamprey and zebrafish genome, we show that an enhancer located *∼* 16 kb downstream of the lamprey *SoxE* gene is able to drive tissue specific reporter expression in the zebrafish neural crest. This contrasts with experiments in the invertebrate chordate, amphioxus, where integration of the entire amphioxus *SoxE* locus and flanking genes into the zebrafish genome resulted in reporter expression in the developing neural tube and tail bud, but not in the neural crest^55^. Together, these results support the hypothesis that the acquisition of novel enhancers in early vertebrates was critical for the evolution of the neural crest GRN. Gain-of-function *cis*-regulatory changes, such as the appearance of new transcription factor binding sites, likely facilitated co-option of preexisting gene batteries, including the pro-chondrocytic *SoxE* genes and other mesenchymal gene programs, into neural crest-like cells at the neural plate border^2^. Indeed, we show that the lamprey *SoxE1* enhancer harbours conserved binding site motifs for several important neural crest transcription factors, including Tfap and SoxE factors, which are known to activate and maintain *SoxE* transcription in the chick^56,57^, zebrafish^58^ and lamprey^29^. HoxA2/B3 sites are also present and possibly control the activity pattern of the enhancer confined to specific regions of cranial neural crest conserved across vertebrate taxa^43^. Our results suggest that the evolution of a combination of key transcription factor binding site motifs was central to neural crest GRN evolution. Conservation of these short motif sequences, without necessarily maintaining their relative position, intervening sequences or exact genomic location, is sufficient to facilitate transcription factor binding and activation of target genes.

## Conclusion

By taking advantage of our highly contiguous germline genome assembly representing lamprey genomic material pre-DNA elimination^17^, we have presented a genome-wide representation of gene expression and chromatin dynamics during lamprey cranial neural crest development. Our analysis of chromatin accessibility across developmental time identifies tissue-specific *cis*-regulatory regions that act as enhancers both in the developing neural tube and the migrating cranial neural crest. Furthermore, by combining ATAC-seq and RNA-seq datasets, we have identified putative non-coding RNA species that are associated with *cis*-regulatory elements in the lamprey. Taken together, our analyses reveal how interrogations of these datasets can uncover critical components of the neural crest GRN that are shared across vertebrates, as well as expose new players whose further investigation will expand our current view of the genetics of neural crest development.

## Methods

### Lamprey husbandry and embryo dissections

Adult sea lamprey (*Petromyzon marinus*) were supplied by the US Fish and Wildlife Service and Department of the Interior. Embryos obtained by *in vitro* fertilisation, were grown to the desired stage as previously described^59^ in compliance with California Institute of Technology Institutional Animal Care and Use Committee protocol #1436. Brook lamprey (*Lampetra planeri*) embryos were collected from a shallow river in the New Forest National Park, United Kingdom, with permission from the Forestry Commission and maintained in filtered river water at 13^*°*^-19^*°*^C. Prior to dissection, embryos were dechorionated in 0.1x Marc’s Modified Ringers buffer (MMR) in a dish lined with 1% agarose. T18, T20 and T21 dorsal neural tubes including premigratory, migrating and/or post-migratory neural crest cells were dissected from the head using an eye-lash knife. T20 and T23 heads were dissected using forceps.

### RNA extraction and library preparation

RNA was extracted from groups of at least 30 dissected dorsal neural tubes at each stage, as well as from whole heads (2 groups of 20) and whole embryos (2 groups of 10) at T20. Tissue was lysed in the Ambion RNaqueous Total RNA Isolation kit lysis buffer (AM1931), set on ice for 15 minutes with occasional vortexing, flash frozen in liquid nitrogen and stored at −80^*°*^C. RNA was extracted using the Ambion RNAqueous Micro Total RNA isolation kit and assessed using the Agilent Bioanalyser. Sequencing libraries were prepared from 100 ng RNA per sample using the NEBNExt Ultra Directional RNA Library Prep Kit for Illumina (E7420) in combination with the NEBNext Poly(A) mRNA Magnetic Isolation Module (E7490) and NEBNext High-Fidelity 2X PCR Master Mix (M0451S). Libraries were indexed and enriched by 15 cycles of amplification. Library preparation was assessed using the Agilent TapeStation and libraries quantified by Qubit. The concentration of library pools was assessed with the KAPA Library Quantification Kit (KK4835). Multiplexed library pools were sequenced using paired-end 75 −100 bp runs on the Illumina NextSeq500 platform for dorsal neural tube libraries and on the Illumina HiSeq2500 platform for T20 heads and embryos.

### ATAC and library preparation

Groups of dissected tissue were collected into L-15 medium (Lifetech) with 10% fetal bovine serum at 19^*°*^C. Tissue was first dissociated in dispase (1.5 mg/ml in DMEM; 10mM Hepes, pH 7.5), followed by the addition of an equal volume of trypsin (0.05% Trypsin; 0.53mM EDTA in HBSS) at room temperature for a total of up to 15 minutes. Dissociated cells were passed over a 40 µm cell strainer into Hanks’ solution (1xHBSS; 10mMHepes; 0.25%BSA) and centrifuged at 500 x g for 7 minutes at room temperature. The supernatant was removed and fresh Hank’s solution applied. 50,000 cells were counted out and centrifuged for 5 minutes at 500 g at 4^*°*^C and washed with cold 2/3 phosphate buffered saline (PBS) by centrifugation for 5 minutes at 500 g at 4^*°*^C. The cells were lysed (10mM Tris-HCl, pH7.4; 10mM NaCl; 3mM MgCl_2_; 0.1% Igepal) and tagmented using the Illumina Nextera kit (FC-121-1030) for 30 minutes at 37^*°*^C as previously described^60^, with the addition of a tagmentation-stop step by the addition of EDTA to final concentration of 50 nM and incubation at 50^*°*^C for 30 minutes. Tagmented DNA was amplified using the NEB Q5 High-Fidelity 2X Master Mix (M0492S) for 14 cycles. Tagmentation efficiency was assessed using Agilent TapeStation and libraries quantified by Qubit. The concentration of ATAC library pools was assessed with the KAPA Library Quantification Kit (KK4835). Multiplexed library pools were sequenced using paired-end 40 bp runs on the Illumina NextSeq500^®^ platform. The high correlation of the mapped ATAC-seq signal between biological replicates at each stage (Pearson’s R*>*0.9) confirms the reproducibility of our experimental approach (Supplementary Fig. 4).

### Pre-processing of Next Generation Sequencing reads

Read quality was evaluated using FastQC^61^. Reads were trimmed to remove low quality bases using Sickle^62^ using the parameters −l 30 −q 20.*RNA-seq analysis* Reads were mapped to the sea lamprey germline genome assembly^19^ using STAR (v2.4.2)^63^ (STAR – genomeDir $GENOME –readFilesIn $R1.fastq $R2.fastq –runThreadN 4 –outFileNamePrefix $PREFIX – readFilesCommand zcat –outSAMstrandField intronMotif –alignEndsType EndToEnd –outReadsUnmapped Fastx –outSAMtype BAM SortedByCoordinate). Separate transcriptomes for dorsal neural tube sample datasets or head and embryo sample datasets were assembled *de novo* with Cufflinks followed by Cuffmerge using default parameters^64^ to make a consensus transcriptome from all the datasets. Read counts for dorsal neural tube datasets were obtained with Subread featureCounts (v1.4.6-p4)^65^ using the Cuffmerge consensus transcriptome in SAF format as a reference (featureCounts -p -B -M -F SAF -s 2 -T 4 -a $SAF -o $OUT $IN.bam).

Differential expression and principal component analysis were performed on the dorsal neural tube read count datasets using DESeq2 (v.1.8.2)^66^. Weighted correlation network analysis (WGCNA)^30^ was performed on the variance stabilised normalised gene count tables generated by DESeq2 according to the pipeline detailed in the online WGCNA tutorial (https://labs.genetics.ucla.edu/horvath/CoexpressionNetwork/Rpackages/WGCNA/). Both of these analyses were run on the R platform (v3.2.1)^67^. The Average normalised gene counts that were associated with ATAC-seq peak-set clusters for stage T21 samples (see ATAC-seq analysis) were plotted in R using ggPlot geom-violin^68^. Output transcript lists from the differential expression analysis and WGCNA were annotated using the gene models associated with the sea lamprey germline genome assembly. *Hox* and *Sox* genes were manually annotated with lamprey-specific gene names. Heatmaps of the average variance stabilised normalised gene counts were generated in R using pheatmap. Gene Ontology (GO) analysis was performed on annotated differentially expressed gene sets using the PANTHER Overrepresentation Test (v11)^69^ with complete GO term databases for *Mus musculus*. Output GO terms were were filtered to only contain terms that were enriched by at least three-fold. Remaining GO terms were summarized with REVIGO^70^.

To identify putative lncRNAs in our transcriptome, first transcripts that overlapped with coding genes in the germline genome annotation on the same strand were eliminated using bedtools(v.2.15.0)^71^ intersect. The remaining transcripts were used in a blastx^72^ search using default parameters against the UniProt/Swiss-Prot database^73^. Any transcripts that shared *>*30% sequence identity with known proteins with an e-value *>*1E-2 were eliminated. Any unspliced transcripts were removed and, using bedtools intersect, the list of putative lncRNAs was limited to transcripts that were within 5kb of a coding gene and originated from the opposite strand to this closest gene. Subread featureCounts was used to determine the length of the remaining transcripts (featureCounts -p -B -F SAF -s 2 -T 4 -a $SAF -o $OUT $IN.bam), and those *<*200 bp in length were eliminated.

### ATAC-seq analysis

Reads were mapped to the sea lamprey germline genome assembly16 using Bowtie2^74^ (bowtie2 –phred33 -p 4 -X 2000 –very-sensitive -x $GENOME −1 $R1.fastq −2 $R2.fastq -S $OUT.sam). Duplicates were removed with Picard (v1.83) MarkDuplicates feature and the distribution of fragment sizes assessed with Picard (v1.83) CollectInsertSizeMetrics^75^ feature. Replicate bam files for each developmental stage were merged with SAMtools^76^ and filtered with BamTools^77^ to remove unpaired reads and reads mapped to the mitochondrial chromosome. Filtered bam files were down-sampled to match the file with the lowest number of reads using Picard (v1.83) DownsampleSam. Down-sampled bam files were sorted by name using SAMtools and paired-end bed files were obtained using bedtools(v.2.15.0)^71^ bamtobed bedpe. Reads were extended to a read length of 100bp. Peak-calling was performed using MACS2^78^ (macs2 callpeak -t $IN.bed f BED name $IN.macs2 –outdir $OUT –shiftsize=100 –nomodel –slocal 1000). Output peak files (.xls) for each developmental stage were converted to bed format and merged with bedtools merge to create one consensus peak set. The consensus peak set was annotated with HOMER (v4.7) annotatePeaks.pl^79^ using the Cuffmerge gene models (with genes less than 1500 bp in length removed) as a reference. Annotated peaks were separated into intergenic, intronic and promoter peak-sets according to their HOMER annotation. Promoter peaks were filtered with bedtools flank to only include peaks that overlapped with a region up to 2 kb upstream of the sea lamprey germline genome gene models (bedtools flank -i promoters.bed -g germline genome.chrom.sizes −l 2000 -r 0 s). Intergenic peaks that overlapped with promoters annotated in the sea lamprey germline genome gene models (i.e. gene models that were not present in the *de novo* cuffmerge assembly) were identified with bedtools intersect and moved to the promoter peak-set. The intergenic and intronic peak-sets were further filtered to only contain peaks that were *<*50,000 bp away from genes that were enriched at stage T21 in comparison to T18 (see RNA-seq analysis). *K* -means clustering of ATAC-seq signal over the final peak-sets was carried out using SeqMINER software^80^ (+/-1500 bp window; no auto-turning; wiggle step: 15; *k*-means enrichment linear). Read counts for the ATAC-seq signal were obtained with Subread featureCounts (v1.4.6-p4) using the peak-set clusters in SAF format as a reference (featureCounts -p -F SAF -T 4 -a peaksetCluster.saf). Correlation analysis on ATAC-seq read count data was performed in R using plot and cor (method=“pearson”). Gene Ontology (GO) analysis was performed on the differentially expressed genes associated with intergenic and intronic “EMT” clusters using the PANTHER Overrepresentation Test (v11)^69^ with complete GO term databases for *Mus musculus*. Output GO terms were filtered to only contain terms that were enriched by at least 1.8-fold. Remaining GO terms were summarized with REVIGO^70^ and subsequently filtered to only contain terms with -log10pvalue less than −1.5. Motif analysis was performed on the intergenic and intronic peak-set clusters with HOMER (v4.7) findMotif.pl. Heatmaps were generated in R using ggPlot geom-tile^68^.

### In vivo enhancer-reporter assays in lamprey

Putative enhancers were amplified by PCR from sea lamprey genomic DNA using primers designed with SnapGene (Clontech), cloned into the HLC GFP reporter vector^41^ by In-Fusion HD cloning (Clontech) and sequenced. ISce-I meganuclease-mediated transgenesis was performed in sea lamprey embryos as described previously^41,43^. At 2-6 hours post fertilisation, single-cell embryos were injected with the ISce-I vector digestion mix at 20 ng/µl and maintained at 18^*°*^C in 0.1x MMR for the remainder of their development. At 1 dpf embryos were transferred to 96-well plates until 6 dpf when they were returned to petri dishes, and screened daily for reporter expression. Live embryos were imaged on a depression slide using a Zeiss Scope.A1 microscope fitted with a Zeiss AxioCam MRm camera and Zeiss ZEN 2012 software (blue edition).

### Cryosectioning and immunostaining

Embryos were fixed at 4^*°*^C overnight in 4% paraformaldehyde in PBS. Fixed embryos were incubated in PBS with 5% sucrose for 4 hours at room temperature, followed by incubation overnight at 4^*°*^C in 15% sucrose in PBS. Embryos were transferred into pre-warmed 7.5% gelatine in 15% sucrose in PBS and incubated overnight at 37^*°*^C, before being transferred to pre-warmed 20% gelatine in PBS. Embryos were embedded in rubber moulds and frozen by immersion in liquid nitrogen. Blocks were cryosectioned at 6-10 µm. Gelatine was removed from the slides by a 5-minute incubation in PBS pre-warmed to 37^*°*^C. Immunostaining was performed as described^81^ using an Alexa-488 conjugated anti-GFP antibody (Rabbit, 1:250; Life Technologies; A21311). Sections were imaged on a Zeiss LSM 780 inverted confocal microscope with Zeiss ZEN 2011 (black edition).

### Zebrafish Husbandry and creation of transgenic lines

This study was carried out in accordance to procedures authorized by the UK Home Office in accordance with UK law (Animals [Scientific Procedures] Act 1986) and the recommendations in the Guide for the Care and Use of Laboratory Animals. Adult fish were maintained as described previously^82^. The lamprey *SoxE1* enhancer was cloned into the Ac/Ds-E1b-eGFP vector (http://www.addgene.org/102417/) using In-fusion cloning (Clontech) and co-injected with Ac transposase mRNA into one-cell-stage zebrafish embryos. Injected F_0_s were screened for founders. Positive F_1_s were grown to reproductive age and backcrossed to F_0_s to obtain embryos with bright reporter expression.

### Whole mount immunostaining

Zebrafish embryos were fixed in 4% paraformaldehyde for 1 hour at room temperature and washed in PBT (1x PBS containing 0.5% Triton X-100 and 2% DMSO). When necessary, embryos were bleached prior to being blocked in 10% Donkey serum in PBT for 2 hours and washed in antibody solution (Rabbit anti-GFP; 1:200 in block; Torrey Pines Cat#TP401) overnight at 4^*°*^C. Embryos were washed several times in PBT before adding the secondary antibody (1:200; Alexa 488 donkey anti-rabbit; ThermoFisher Scientific; A21206) in combination with Hoescht (1:1000) for two hours at room temperature. After several PBT washes, embryos were imaged on a Zeiss LSM 780 upright confocal microscope with Zeiss ZEN 2011 (black edition).

### Splinkerette PCR

Splinkerette analysis was performed as previously described^83,84^. Five positive F_2_ zebrafish embryos from a single F_1_ parent outcrossed to wild type were collected and genomic DNA extracted. Approximately 500 ng of genomic DNA was digested overnight with AluI in a 30 µl reaction. Digested genomic DNA was purified using phenol-chloroform followed by ethanol precipitation before ligation with annealed splink-erette adaptors (CGAATCGTAACCGTTCGTACGAGAATTCGTACGAGAATCGCTGTCCTCTCCGGC- CACAGGCGATTAT and ATAATCGCCTGTGGCCAAATCTATACGTATAGAT) using T4 DNA ligase at 16^*°*^C overnight in a thermal cycler. The adaptor-ligated genomic DNA was purified using Zymo Research Clean & Concentrate (Cat.#D4003) and 20 ng of purified product used in a primary PCR reaction. PCR was performed using the following primers: CGAATCGTAACCGTTCGTACGAGAA (binding to adaptor) and GTTTCCGTCCCGCAAGTTAA (binding to Ds-5^*′*^ integration arm), with 63^*°*^C annealing temperature and 3 mins extension time. 1 µl of primary PCR reaction was then used in 50 µl nested PCR reaction using the following primers: TCGTACGAGAATCGCTGTCCTCTC (binding to adaptor) and CGGTA-GAGGTATTTTACCGAC (binding to Ds-3^*′*^ integration arm), with 60^*°*^C annealing temperature and 5 mins extension time. The nested PCR was run on agarose gel to visualise number of integrations.

## Acknowledgements

We thank S. Shimeld for access to brook lamprey embryos and H. Parker for the lamprey HLC vector. This work was supported by a Leverhulme Research Grant to T.S.S. (RPG-2015-026), the National Institute of General Medical Sciences of the National Institutes of Health grants to J.J.S. (R01GM104123) and C.T.A. (R24GM095471), a Wellcome Trust Institutional Strategic Support Fund grant (H2RZKC00) to T.S.S. and D.H., a Junior Research Fellowship (Trinity College, Oxford), the Sydney Brenner Fellowship, a Company of Biologists Travelling Fellowship (DEVTF-150403) and an EMBO Short Term Fellowship to D.H., and a Clarendon Fund Fellowship to V.C.M.

## Author contributions

D.H. and T.S.S. conceived this research programme. D.H. collected RNA-seq and ATAC-seq data, performed and analysed lamprey reporter expression assays and bioinformatics analysis. V.C.-M. performed zebrafish transgenesis, splinkerette assay and immunostaining. D.G. assisted on the analysis of RNA-seq and ATAC-seq data. S.G. and M.E.B. assisted with access to sea lamprey embryos. J.S. and C.T.A. provided access to the draft sea lamprey germline genome assembly. D.H. and T.S.S. discussed ideas and interpretations and wrote the manuscript. D.H., M.E.B., and T.S.S. edited the manuscript and all authors commented on it.T.S.S. supervised the study.

---

**Figure S1.**
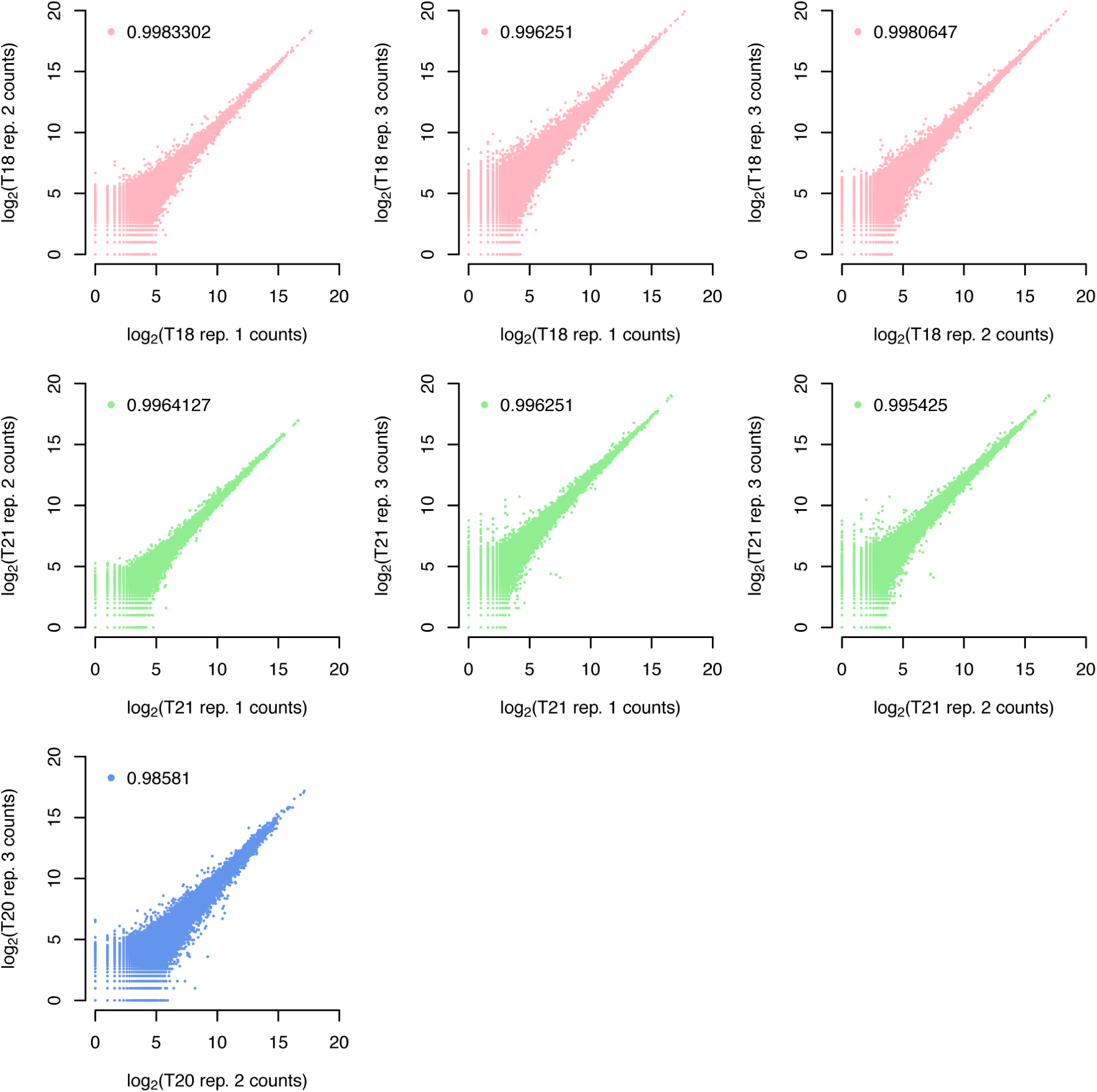
Reproducibility of RNA-seq experiments. Scatterplots between replicate dorsal neural tube RNA-seq datasets at Tl8, T20 and T21 shows replicates are highly correlated. Pearson correlation coefficients (r) for all comparisons are given.

**Figure S2.**
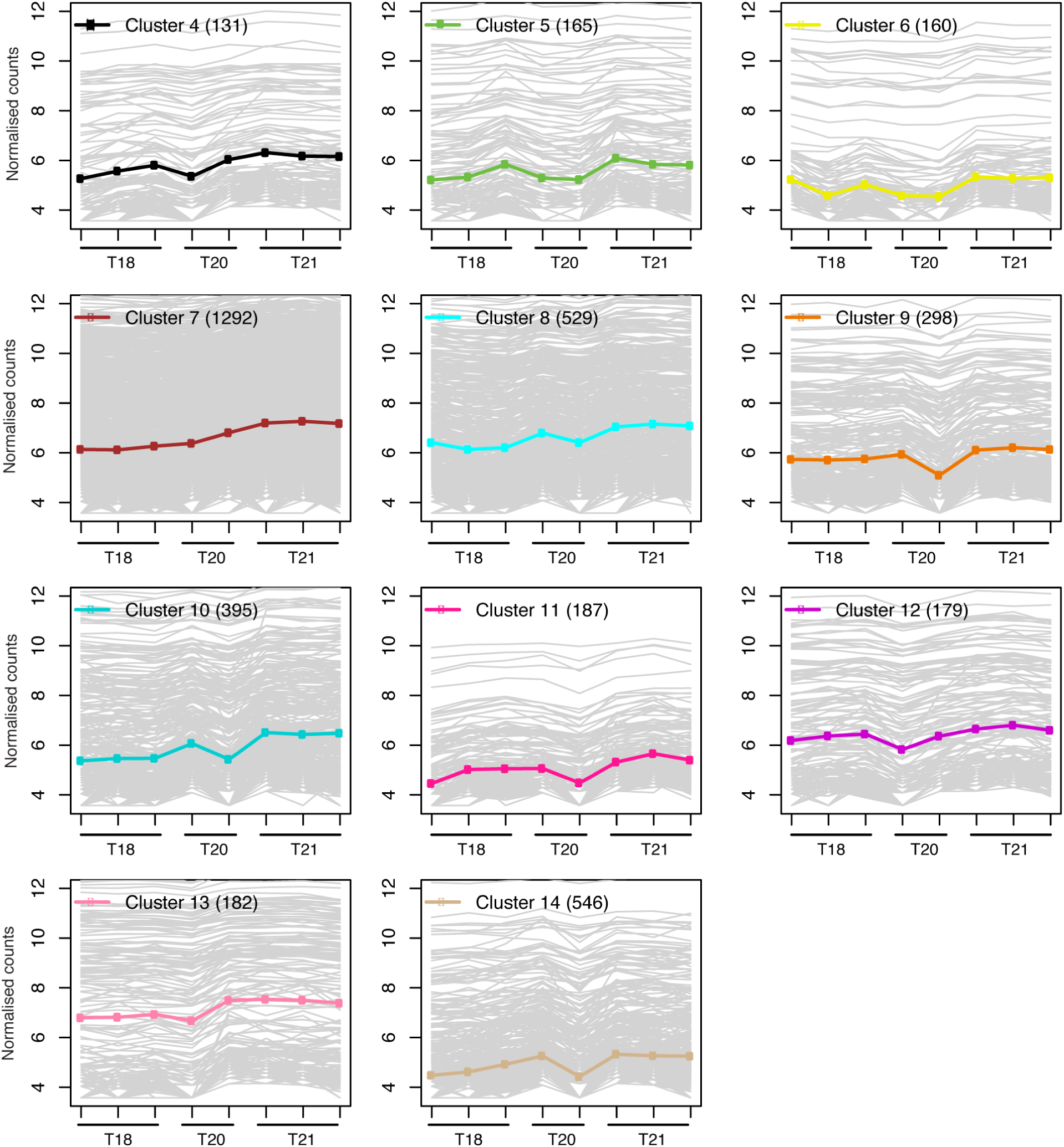
Additional WGCNA clusters showing significantly higher gene expression at T21.

**Figure S3.**
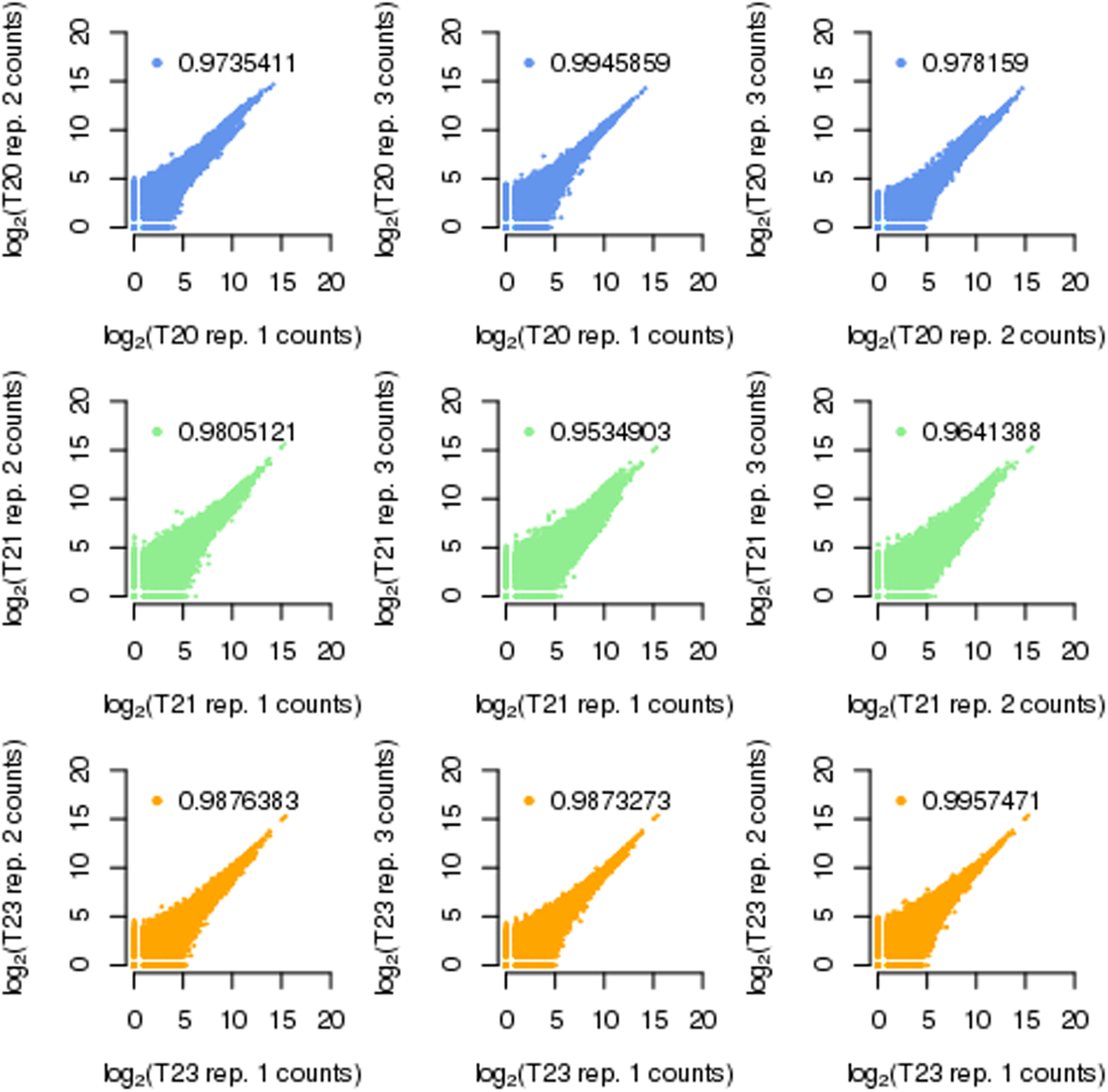
Reproducibility of ATAC-seq experiments. Scatterplots between replicate dorsal neural tube (T20 and T21) and head (T23) ATAC-seq datasets shows replicates are highly correlated. Pearson correlation coefficients (r) for all comparisons are given.

**Figure S4.**
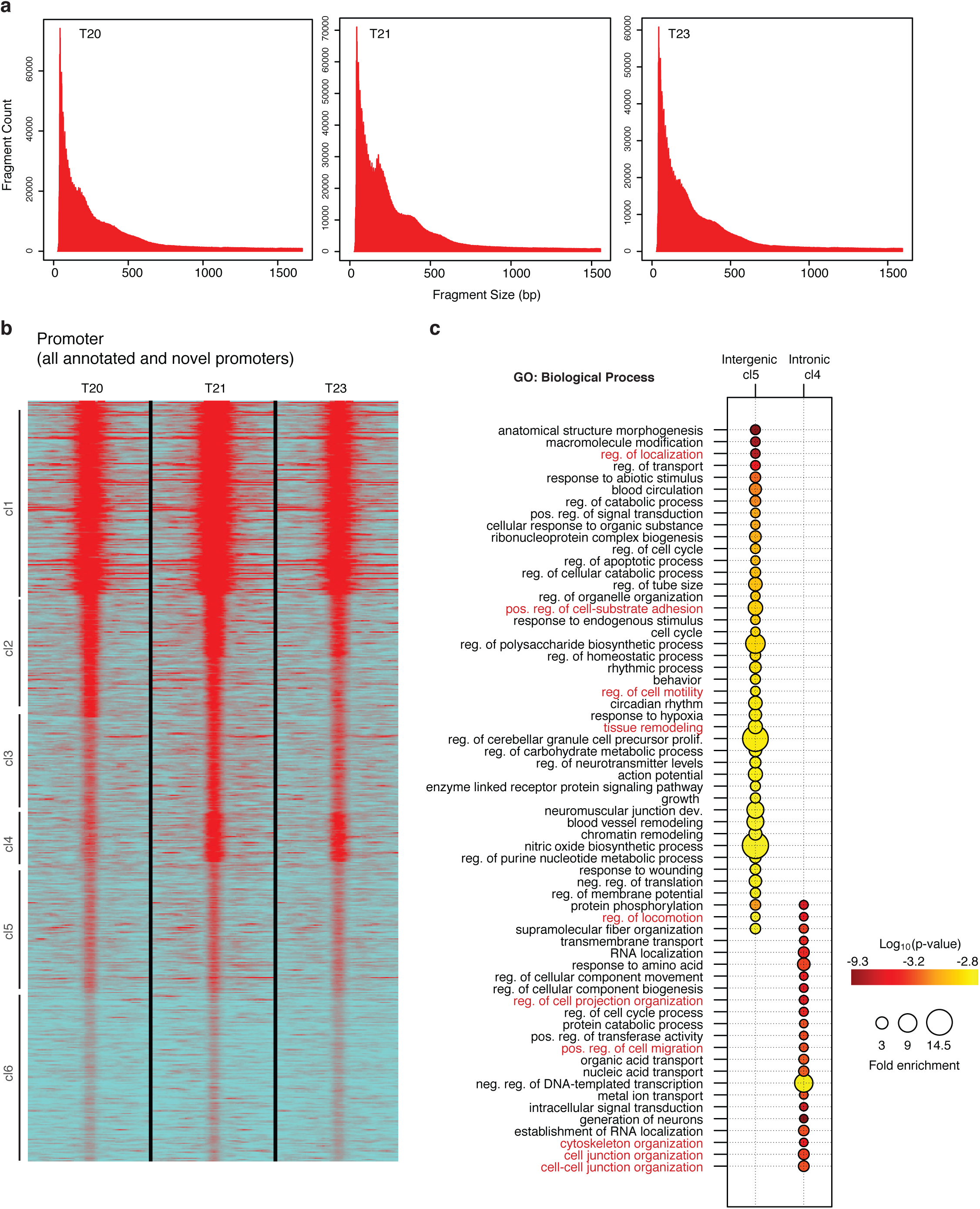
ATAC-seq analysis. **a,** Histograms of fragments size for representative ATAC-seq samples at T20, T21 and T23 shows a periodicity of ∼150 bp, corresponding to nucleosome protected fragments. **b,** Heatmap depicting k-means linear enrichment clustering of the promoter peakset (annotated and novel promoters) associated with the “Promoter” violin plot shown in Figure 2e. **c,** Bubble plots summarizing enrichment and p-values for the most significant GO biological process terms for the differentially expressed genes associated with Intergenic peaks k-means cluster 5 and Intronic peaks k-means cluster 4 (‘EMT’ clusters, see Fig. 2d). GO terms associated with cell migration are highlighted in red. Values shown are for terms that were more than 1.8-fold enriched.

**Figure S5.**
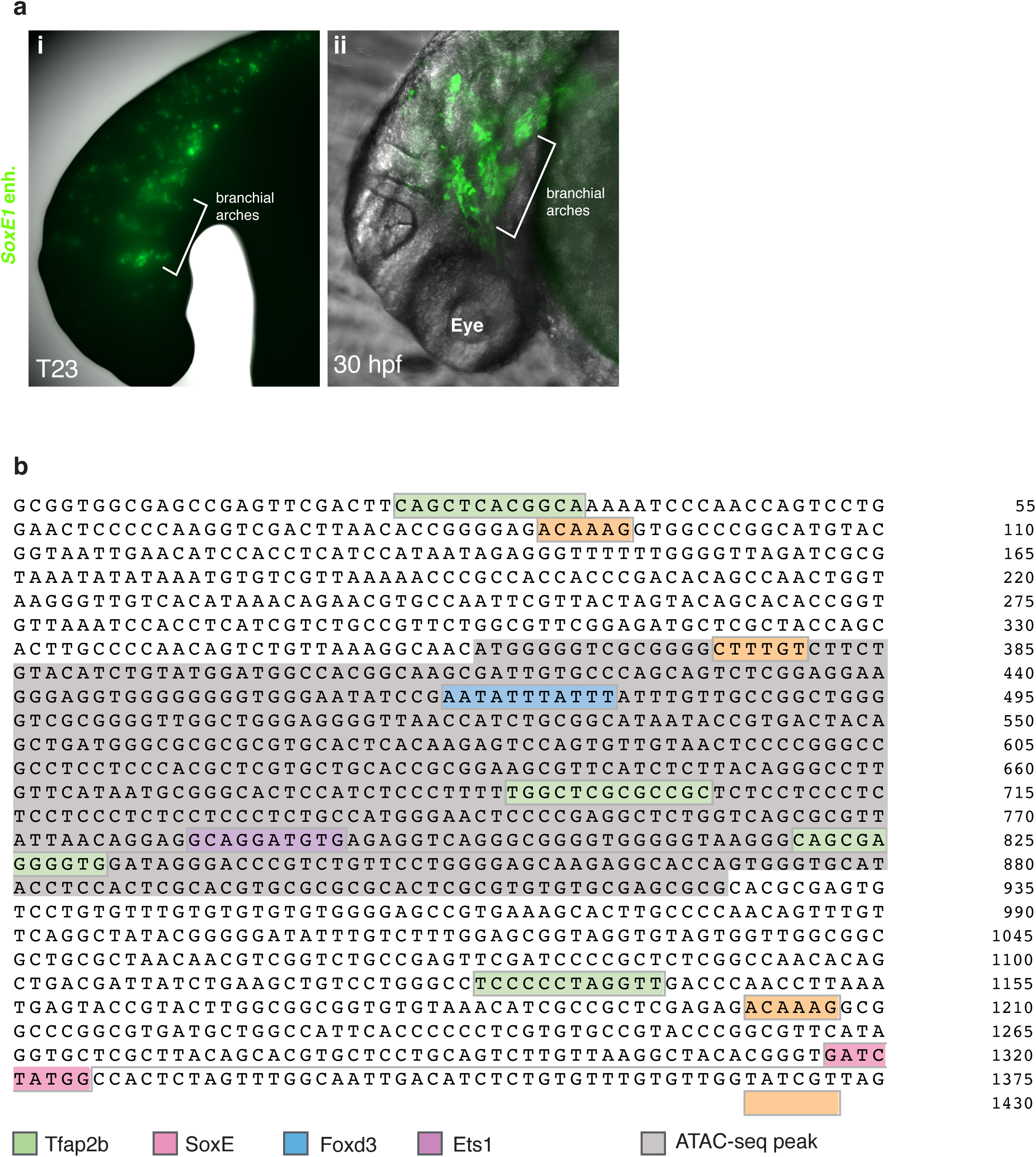
Sea lamprey *SoxE1* enhancer activity is conserved in gnathostomes. **a,** High magnification views of *SoxE1* enhancer-reporter expression in equivalently staged lamprey (T23, i) and zebrafish (30 hpf, ii) embryos showing GFP expression in the developing branchial arches. **b,** The lamprey *SoxE1* enhancer sequence with putative transcription factor binding motifs indicated by coloured boxes.

